# Structural characterization of a pseudaminic acid-modified lateral flagellar filament from *Vibrio alginolyticus*

**DOI:** 10.64898/2026.09.01.748753

**Authors:** Miki Kinoshita, Tohru Minamino, Hajime Nakatani, Atsuo Suzuki, Toshifumi Takao, Emi Mishiro-Sato, Seiji Kojima, Keiichi Namba, Michio Homma

## Abstract

Bacterial flagellar filaments are often modified by glycans, but the structural basis and physiological significance of flagellin glycosylation remain poorly understood in many species. *Vibrio alginolyticus* produces lateral flagella for surface-associated motility in viscous environments, with LafA forming its filament as flagellin. A *maf* homolog encoding a putative flagellin glycosylation factor is located immediately downstream of *lafA*, suggesting that the lateral filament is glycosylated. To investigate this possibility, we purified the lateral flagellar filament from *V. alginolyticus* and determined the structure at 2.37 Å resolution by electron cryomicroscopy. Upon model building of LafA in the map, we identified additional densities connected to five serine residues, possibly corresponding to O-linked pseudaminic acid modifications. Mass spectrometric analyses identified these modifications as pseudaminic acid attached to Ser148, Ser173, Ser183, Ser189, and Ser197. Deletion of *maf* abolished lateral flagella formation and motility, and these defects were restored by complementation with *maf*. These results demonstrate that the *Vibrio* lateral flagellar filament is extensively modified by pseudaminic acid and that Maf is required for filament formation. Our findings provide the first structural insight into the glycosylation of *Vibrio* lateral flagellar filament and establish a framework for understanding the role of flagellin glycosylation in surface-associated motility.

**IMPORTANCE:** A gene encoding a putative flagellin glycosylation factor (Maf) is located immediately downstream of *lafA*, the gene encoding lateral flagellin of *Vibrio alginolyticus*, suggesting that the lateral flagellar filament is glycosylated. Guided by this assumption, we determined the structure of the lateral flagellar filament by electron cryomicroscopy with mass spectroscopy and identified five pseudaminic acid modifications on the filament surface. Genetic analyses further demonstrated that Maf is required for lateral flagellar formation and motility. These results provide the first structural characterization of a glycosylated *Vibrio* lateral flagellar filament and reveal a close link between flagellin glycosylation and motility organelle assembly.

## INTRODUCTION

Protein glycosylation is a widespread post-translational modification that occurs in all domains of life (1, 2). In bacteria, both N-linked and O-linked glycosylation pathways have been identified, and numerous cell-surface proteins are known to be modified by diverse glycans. Among these proteins, flagellins are one of the best-characterized substrates for O-linked glycosylation (1, 3, 4).

The bacterial flagellum is a rotary organelle for motility consisting of the basal body, hook, and filament (5, 6). Flagellin is the structural subunit of the flagellar filament. Flagellin glycosylation has been reported in a wide range of Gram-negative and Gram-positive bacteria and contributes to diverse biological processes, including motility, filament assembly, host colonization, and immune evasion (7). Glycosylation sites are typically located on surface-exposed regions of flagellin and are often modified by unusual sugars, including pseudaminic acid and related nonulosonic acids (8–16). Despite increasing knowledge of bacterial glycosylation pathways, the structural basis and physiological significance of flagellin glycosylation remain poorly understood in many bacterial species.

Previous genetic studies have identified motility-associated factor (Maf) proteins as key components required for flagellin glycosylation. Maf homologs are frequently encoded adjacent to flagellin genes, and deletion of *maf* has been shown to abolish flagellin glycosylation in several bacterial species (17–20). Structural studies revealed that Maf proteins contain a central domain structurally related to the *Campylobacter jejuni* sialyltransferase CstII (PDB ID: 1RO7), supporting their proposed role in flagellin glycosylation; however, the molecular relationship between Maf-dependent glycosylation and flagellar filament assembly remains incompletely understood.

*Vibrio alginolyticus* possesses two distinct flagellar systems: a sodium-driven polar flagellum for swimming in liquid environments and proton-driven lateral flagella that promote surface-associated motility under viscous conditions. The structure, assembly, and function of the polar flagellum have been well studied (21–24). In contrast, the functional role of the lateral flagella of this bacterium remains poorly understood. Recent studies have shown that gelatin supplementation markedly enhances lateral-flagella-dependent swarming, highlighting the importance of lateral flagella for surface motility (25). Despite their physiological importance, little is known about the molecular architecture of lateral flagellar filaments. Interestingly, we found that a *maf* homolog is located immediately downstream of *lafA*, which encodes the lateral flagellin protein LafA (Fig. 1), suggesting that the lateral flagellar filament may be glycosylated.

**Fig. 1.**
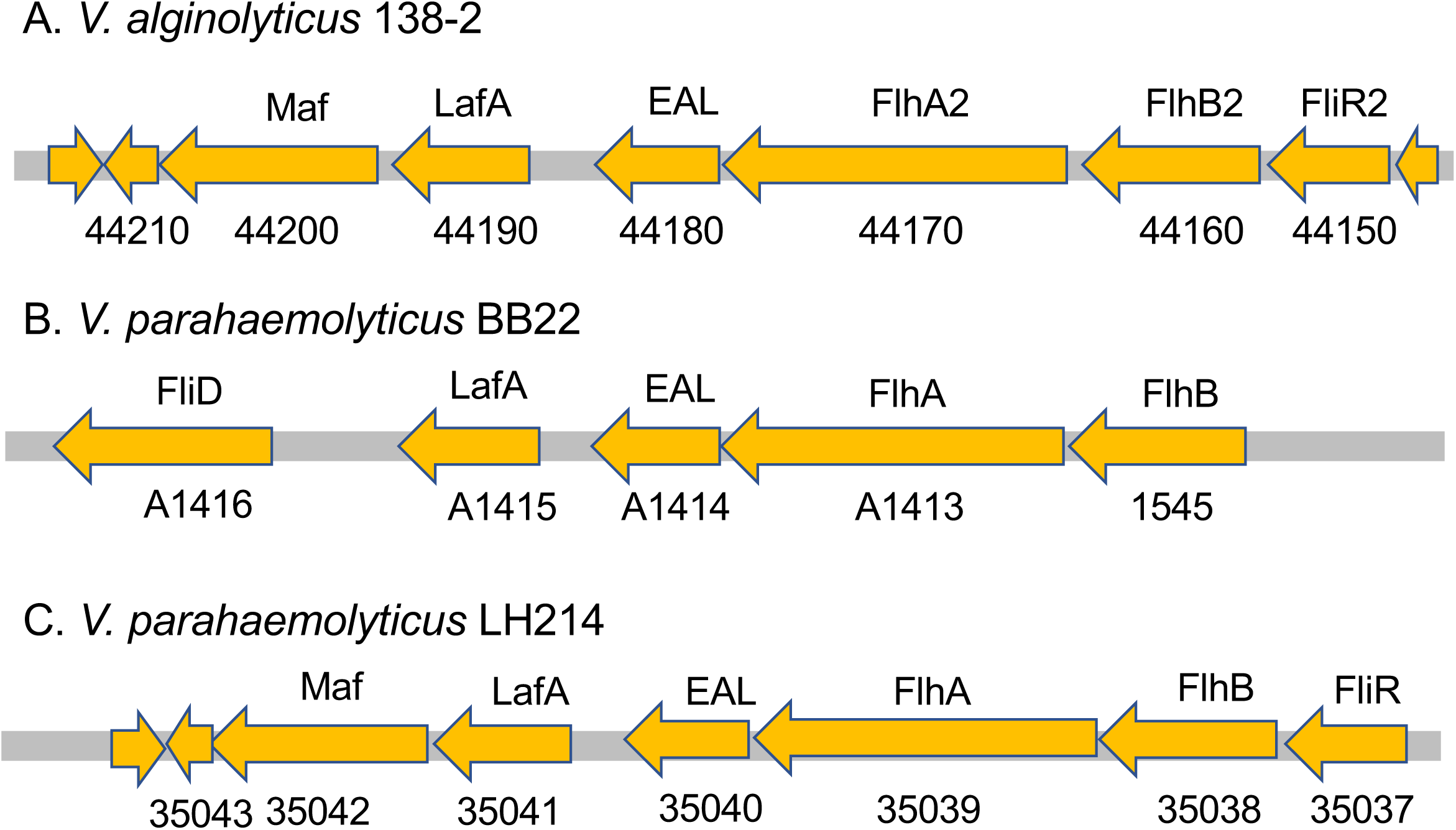
Genetic organization of the *lafA* locus in *Vibrio* species. Genetic organization of the chromosomal region surrounding *lafA* in (A) *V. alginolyticus* 138-2, (B) *V. parahaemolyticus* BB22, and (C) *V. parahaemolyticus* LH214. Predicted gene products are indicated above the arrows, and locus identifiers are shown below. A *maf* homolog is located immediately downstream of *lafA* in *V. alginolyticus* 138-2 and *V. parahaemolyticus* LH214, but is absent from the corresponding locus of *V. parahaemolyticus* BB22.

In this study, using electron cryomicroscopy (cryoEM), we determined the structure of the lateral flagellar filament isolated from *V. alginolyticus* at 2.37 Å resolution. We identified five O-linked pseudaminic acid modifications on the filament surface by combining cryoEM structural analysis with mass spectrometry. Furthermore, genetic analyses demonstrated that Maf is required for lateral flagellar filament formation and motility. These findings provide structural insight into glycosylated lateral flagellar filaments and establish a framework for understanding the role of flagellin glycosylation in *Vibrio* surface-associated motility.

## RESULTS

### Genetic organization of the *laf* and *maf* genes

The lateral flagellin gene *lafA* encodes the structural subunit of the lateral flagellar filament responsible for surface-associated motility of *V. alginolyticus* under viscous conditions. Unlike *Salmonella* FliC, which consists of four domains (D0–D3) (5, 6, 26), LafA contains only the conserved D0 and D1 domains (Fig. S1), leaving the surface-exposed region of domain D1 as a potential site for interactions with the external environment.

To investigate whether LafA may be modified by glycans, we analyzed the genomic region surrounding *lafA* in the *V. alginolyticus* strain 138-2 genome (AP022860). We identified a *maf* homolog immediately downstream of *lafA* (Fig. 1). A similar genetic organization was also found in *Vibrio parahaemolyticus* strain LH214, which is closely related to *V. alginolyticus*. In contrast, neither *V. parahaemolyticus* strain BB22 nor *V. alginolyticus* strain ATCC17749 contained a *maf* homolog adjacent to *lafA* (Fig. 1).

Comparison of LafA amino acid sequences revealed that the D1 domain exhibited relatively low sequence conservation between *Vibrio* strains possessing *maf* and those lacking *maf* (Fig. S2). Because the D1 domain forms the outer surface of the filament, these observations raised the possibility that LafA may undergo Maf-dependent modification in *V. alginolyticus*.

### CryoEM image analysis of the lateral flagellar filament

The lateral flagellar filament was purified from *V. alginolyticus* strain YM19 as described in Material and Methods (Figs. S3–S5) and subjected to cryoEM image analysis (Fig. S6A, B). A total of 6,598 micrographs were selected after CTF-based quality filtering, and helical reconstruction of 566,405 segmented particles yielded a 3D density map at 2.37 Å resolution (EMD-81163) (Figs. 2A and S6C and Table 1), allowing construction of an accurate atomic model of the lateral flagellar filament (PDB ID: 27HA) (Fig. 2B).

**Fig. 2.**
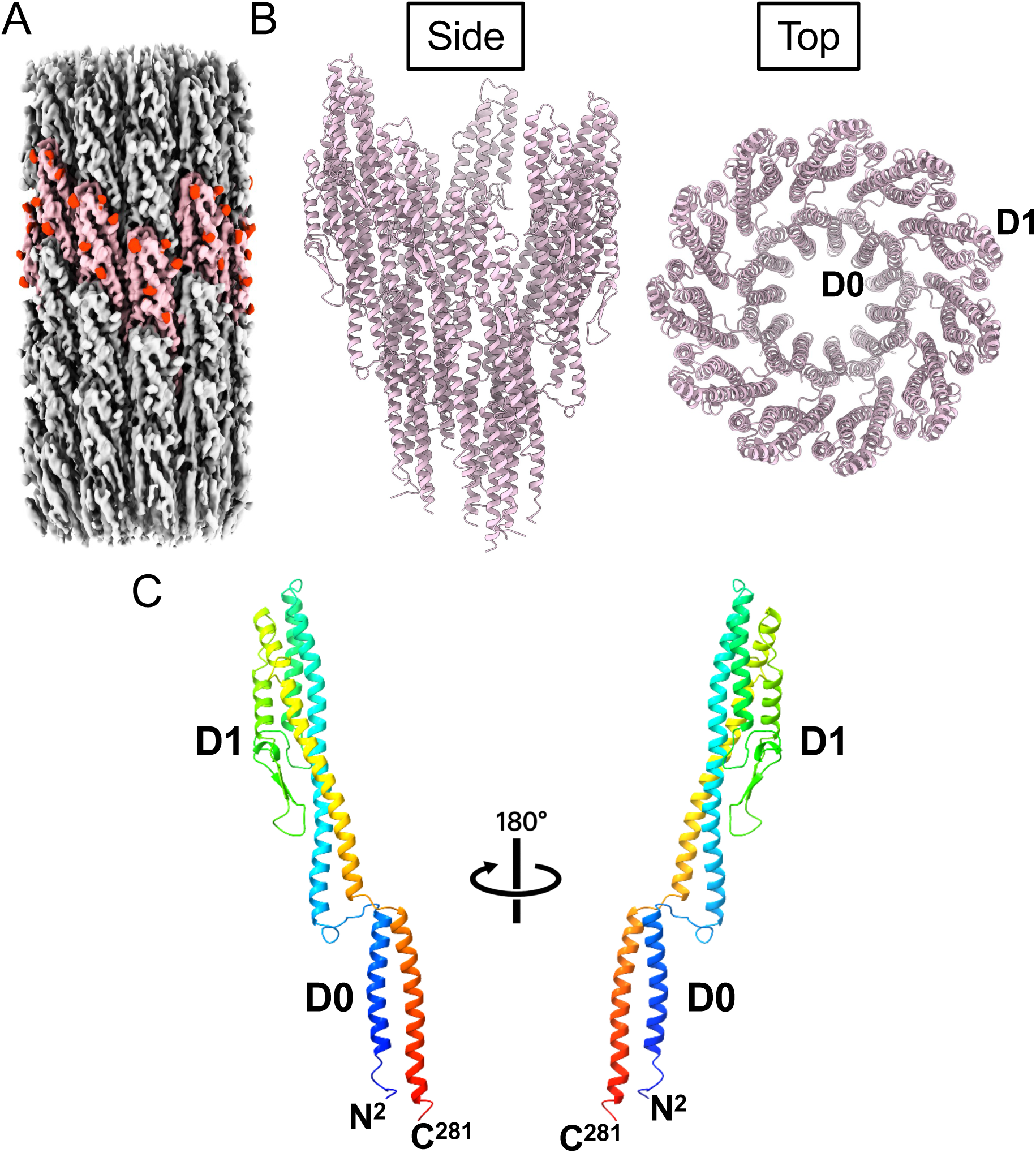
CryoEM structure of *V. alginolyticus* lateral flagellar filament. (A) Side view of the cryoEM density map of the LafA filament (EMD-81163). Small globular densities in red on the surface represent additional densities observed adjacent to five serine residues (Ser148, Ser173, Ser183, Ser189, and Ser197) that are glycosylated. Eleven LafA subunits are colored pink to indicate the subunits in 2 turns of 1-start helix of the helical lattice of the filament. (B) Cα ribbon representation of the atomic model of the filament built into the cryoEM density map (PDB ID: 27HA). The filament comprises 11 protofilaments. Domain D0 forms the inner core of the filament, whereas domain D1 forms the outer surface of the filament. (C) Atomic model of a LafA subunit. The Cα backbone is colored in rainbow from the N-terminus (blue) to the C-terminus (red). Domain D0 is composed of residues Y9–T34 and F245–S278, and domain D1 is composed of residues A45–T241. The N- and C-terminal spoke regions, comprising residues G35–D44 and D242–D244, respectively, connect the D0 and D1 domains.

**Table 1.** CryoEM data collection, processing, and refinement statistics.

|  |  |
| --- | --- |
| Dataset | Glycosylated<br>LafA filament |
| EMDB | EMD-81163 |
| PDB | 27HA |
| **Data collection and processing** |  |
| Magnification | 60,000 |
| Voltage (kV) | 300 |
| Electron exposure (e-/Å<sup>2</sup>) | 80 |
| Defocus range (μm) | -0.7 – -2.2 |
| Pixel size (Å) | 0.786 |
| Symmetry imposed | Helical |
| Imported movies (no.) | 6,825 |
| Initial particle images (no.) | 2,905,795 |
| Final particle images (no.) | 566,405 |
| Map resolution (Å) | 2.37 |
| FSC threshold | 0.143 |
| Helical parameters |  |
| Rise(Å) | 4.728 |
| Twist(°) | 65.351 |
| **Refinement** |  |
| Model resolution | 3.6 |
| FSC threshold | 0.143 |
| **Model composition** |  |
| Non-hydrogen atoms | 23,221 |
| Protein residues | 3080 |
| Ligands | 55 |
| **R.m.s. deviations** |  |
| Bond length (Å) | 0.004 |
| Bond angles (°) | 0.626 |
| **Validation** |  |
| MolProbity score | 1.92 |
| Clash score | 9.54 |
| Rotamer outliers (%) | 3.54 |
| **Ramachandran plot** |  |
| Favored (%) | 98.30 |
| Allowed (%) | 1.70 |
| Disallowed (%) | 0 |

The lateral flagellar filament is composed of LafA, which a small flagellin consisting exclusively of the two conserved domains D0 and D1 (Fig. 2C). Despite lack of the large surface-exposed domains, D2 and D3, the overall architecture of LafA closely resembles those of *Salmonella* flagellins FliC (PDB ID: 1UCU) and FljB (PDB ID: 6JY0). Structural superposition of LafA with FliC and FljB yielded root-mean-square deviations of 0.936 Å and 0.949 Å, respectively, indicating a high degree of structural conservation within the D0 and D1 core domains (Fig. S7).

The LafA atomic model we built and refined in the cryoEM density map revealed five additional densities adjacent to Ser148, Ser173, Ser183, Ser189, and Ser197 on the filament surface (Fig. 3A), suggesting that these residues may carry glycan modifications.

**Fig. 3.**
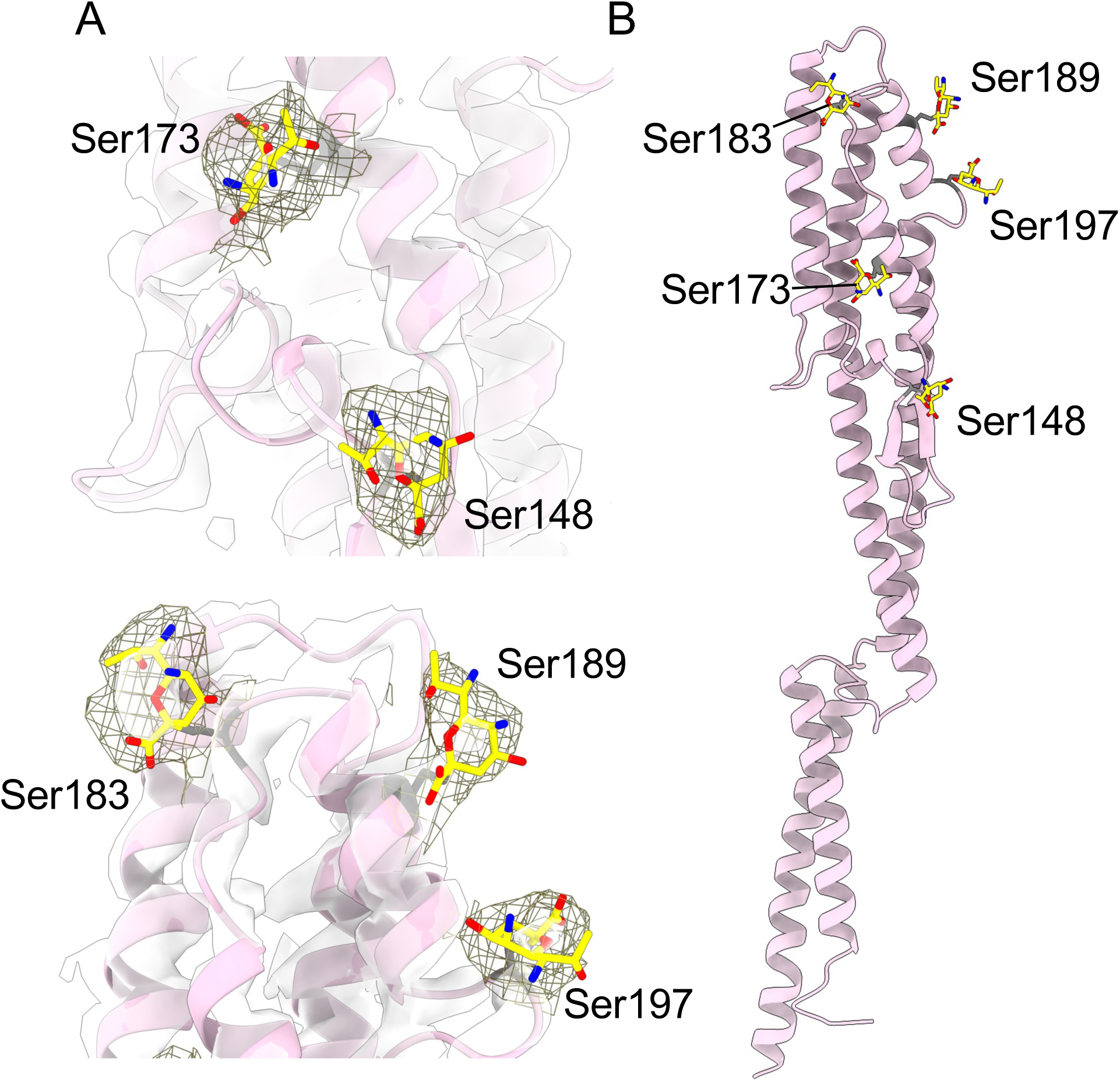
Glycosylation sites identified on the LafA filament. (A) Atomic model of LafA built into the cryoEM density map. Additional densities that not occupied with LafA residues and observed adjacent to five serine residues (Ser148, Ser173, Ser183, Ser189, and Ser197) are shown in a mesh representation. Pseudaminic acid (Pse) models identified by mass spectrometric analysis (Fig. 4) were fitted into the corresponding cryoEM densities and are shown as stick models. (B) Ribbon representation of a glycosylated LafA subunit viewed from the side. The five glycosylated serine residues and their attached Pse moieties are shown as stick models.

### Identification of pseudaminic acid modifications on LafA

To determine the chemical nature of these modifications, we analyzed tryptic peptides of LafA by LC/ESI-MS/MS (Fig. 4A). Most peptides were successfully assigned to the protein sequence (Fig. 4B). However, peaks 10 and 16 remained unassigned; these yielded multiply charged ions with the most intense peaks observed at m/z 1300.5 (2+) and 1332.9 (4+), corresponding to deconvoluted MH^+^ ions of 2600.0 and 5328.6, respectively. These MH^+^ values were 315.9 and 1264.5 Da higher than those of the corresponding unmodified tryptic peptides. Furthermore, the ESI-TOFMS spectrum of peptide 10 exhibited a characteristic fragment ion corresponding to a neutral loss of 316.1 Da from the precursor ion (Fig. 4C). Because pseudaminic acid and related nonulosonic acid derivatives have previously been identified in *Vibrio* flagellins, the observed mass increments of 315.9 and 1264.5 Da were most likely modified with one and four di-*N*-acetylated pseudaminic acid (Pse) derivatives (residue mass of 316.1 Da), respectively. Although MS/MS analysis alone could not pinpoint the exact modification sites, the cryoEM density map clearly assigned the modifications to Ser-148 in the peptide 10, and to S-173, Ser-183, Ser-189, and Ser-197 in the peptide 16.

**Fig. 4.**
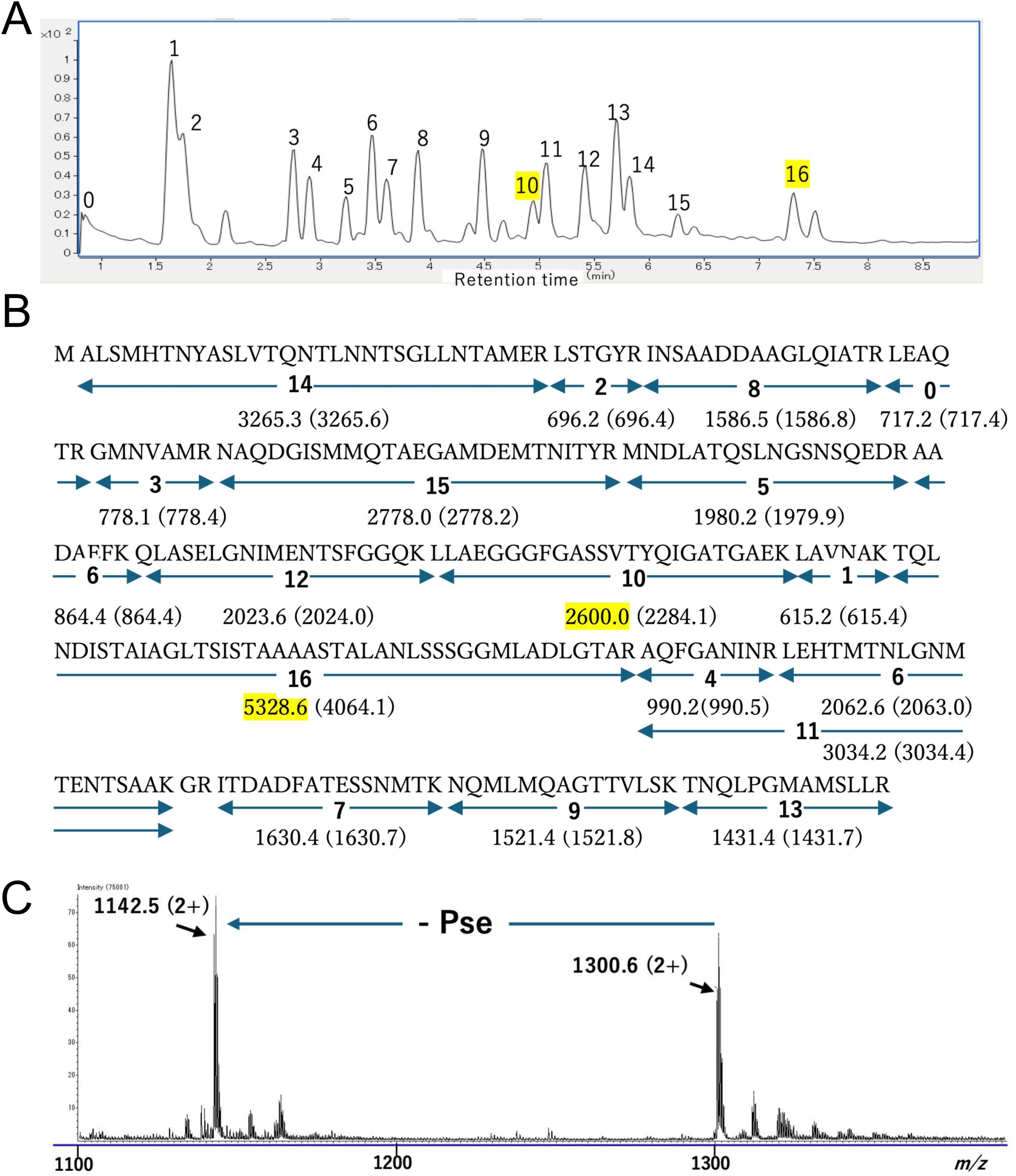
Mass spectrometric identification of pseudaminic acid modifications on LafA. (A) Total ion chromatogram obtained by LC/ESI-MS analysis of tryptic peptides derived from LafA. Peaks 10 and 16 correspond to modified peptides exhibiting mass shifts relative to the predicted unmodified peptides. (B) Sequence coverage map of LafA. Identified tryptic peptides are indicated along the amino acid sequence. Peptides 10 and 16 were assigned as peptides carrying one and four pseudaminic acid (Pse) modifications, respectively. Observed MH+ values and corresponding theoretical masses are shown. (C) ESI-TOFMS spectrum of peptide 10. The precursor ion at m/z 1300.6 (2+) produced a characteristic fragment corresponding to a neutral loss of 316.1 Da, consistent with a di-N-acetylated pseudaminic acid residue.

To further validate the glycan modification, we treated the LafA monomer obtained by heat depolymerization of the flagellar filament with glycosidases and analyzed by SDS-PAGE with CBB staining (Fig. S8). Treatment of LafA with endo-α-N-acetylgalactosaminidase (Endo-α) alone did not result in any changes in the LafA band. However, treatment with a mixture of Endo-α and Sialidase resulted in a change in the CBB staining of the band. No change in the profiles of CBB staining bands was observed when the flagellar filament without heat treatment was treated by these enzymes. These observations are consistent with the presence of sialic-acid-related glycan modifications on LafA.

Taken together, the cryoEM, MS, and glycosidase analyses support the assignment of five pseudaminic acid residues on LafA. Incorporation of five Pse residues into the cryoEM density map produced an excellent fit to the experimental density (Fig. 3A). The five glycosylation sites are distributed approximately evenly over the filament surface, generating a nearly continuous glycosylated outer layer (Fig. 3D and Fig. S9).

### Maf is required for lateral flagellar formation and surface-associated motility

To investigate the role of Maf in lateral flagellar filament formation, we constructed the Δ*maf* mutant of *V. alginolyticus* strain YM19. Because YM19 produces only lateral flagella, motility of this strain directly reflects lateral-flagellum-dependent motility. The Δ*maf* mutant exhibited a severe motility defect in both 0.3% soft agar and 1.25% agar media compared with the parental YM19 strain (Fig. 5A and Fig. S10). Introduction of a plasmid expressing *maf* restored motility of the Δ*maf* mutant (Fig. S10), indicating that the motility defect resulted from loss of the Maf function.

**Fig. 5.**
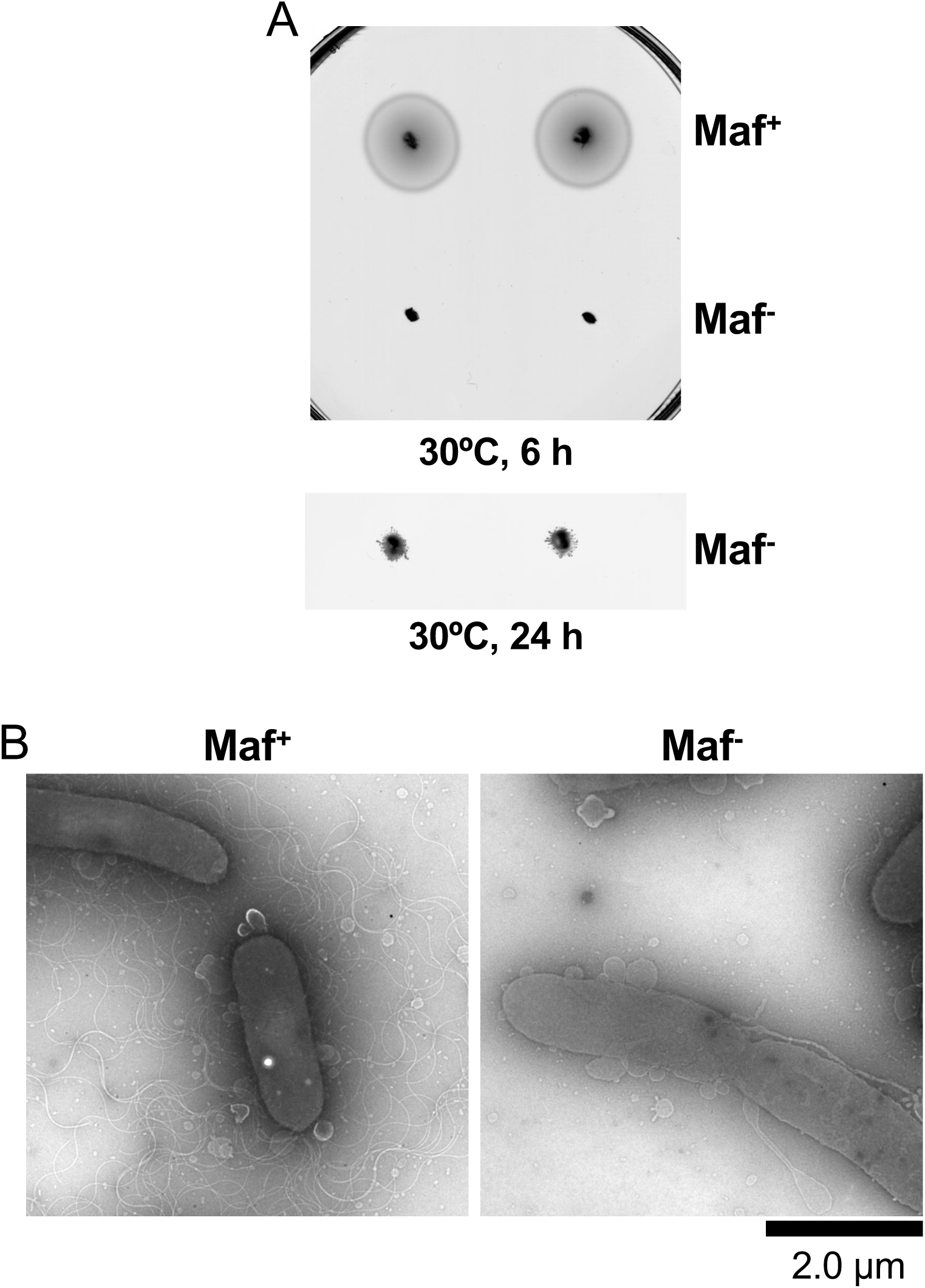
Effect of deletion of *maf* on lateral flagella-dependent motility. (A) Swimming motility assays on soft agar plates. Overnight cultures of *V. alginolyticus* YM19 (Maf^+^) and its Δ*maf* mutant (Maf^−^) were inoculated onto VNG plates containing 0.3% agar and incubated at 30°C. Representative motility halos were recorded after 6 h (upper panel) and 24 h (lower panel). Deletion of *maf* markedly impaired motility compared with the parental strain. (B) Representative negative-stain electron micrographs of *V. alginolyticus* YM19 (Maf^+^) and the Δ*maf* mutant (Maf^-^). Cells were collected from the edges of colonies grown on VNG plates containing 1.25% agar and 1% gelatin, stained with 1% phosphotungstic acid, and observed by transmission electron microscopy. Numerous lateral flagellar filaments are observed on YM19 cells, whereas no lateral flagella are detected on the Δ*maf* mutant.

To determine whether the motility defect was associated with impaired lateral flagellar assembly, cells grown on 1.25% agar plates were examined by negative-stain electron microscopy. Numerous lateral flagella were observed on wild-type YM19 cells, whereas no lateral flagella were detected on the Δ*maf* mutant cells (Fig. 5B).

We next examined extracellular accumulation of LafA. Lateral flagellar fractions were isolated from the three strains indicated in Fig. 6, treated with Triton X-100, and analyzed by SDS-PAGE (Fig. 6). A protein band corresponding to LafA (∼30 kDa) was readily detected in the pellet fraction of the Δ*lafA* mutant complemented with a *lafA* plasmid and the Δ*maf* mutant complemented with a *maf* plasmid (lanes 1 and 2). In contrast, LafA was not detected in the Δ*maf* mutant expressing *lafA* alone (lane 3). These results indicate that expression of LafA itself is insufficient for lateral filament formation in the absence of Maf and that Maf is required for extracellular accumulation of LafA and normal assembly of the lateral flagellar filament.

**Fig. 6.**
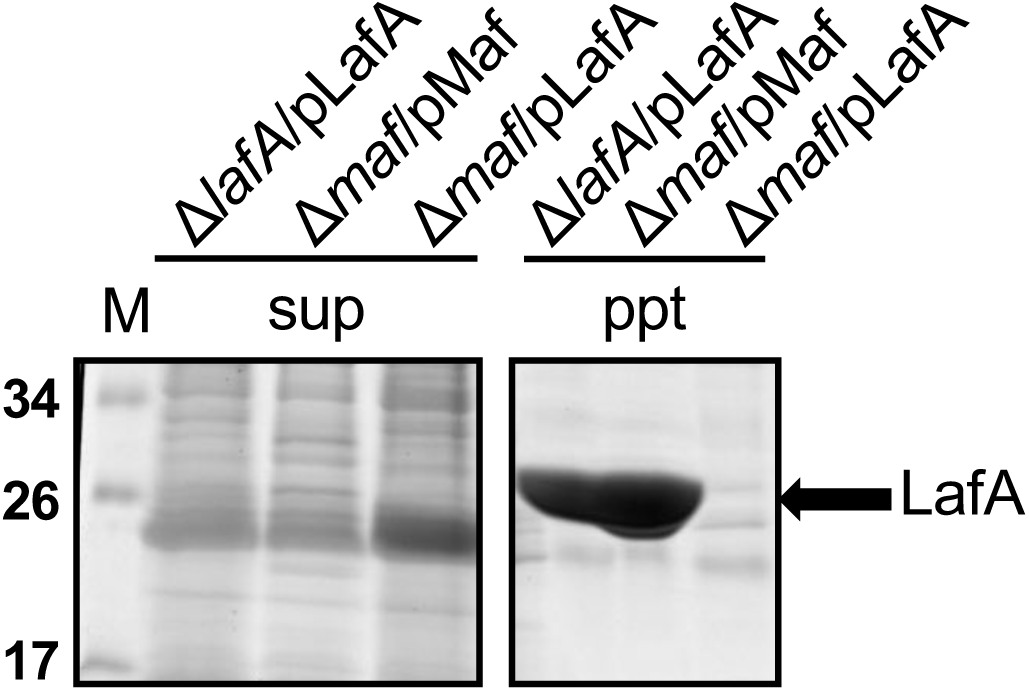
Recovery of extracellular LafA by complementation of the *maf* mutant. Extracellular flagellar fractions were prepared from the strains indicated above the lanes [1, YM19Δ*lafA*/pTSK191 (*lafA*); 2, YM19Δ*maf*/pTSK193 (*maf*); 3, YM19Δ*maf*/pTSK191 (*lafA*)] grown on agar plates and analyzed by SDS-PAGE followed by CBB staining. Samples were separated into supernatant (sup) and pellet (ppt) fractions after Triton X-100 treatment and centrifugation. A protein band corresponding to LafA (∼30 kDa) was detected in the pellet fraction of strains complemented with *maf* or *lafA*, whereas LafA was not detected in the Δ*maf* mutant expressing *lafA* alone. Lane M, molecular weight markers (kDa).

Taken together, these results demonstrate that Maf is required for lateral flagellar formation and is essential for efficient lateral-flagellum-dependent motility in *V. alginolyticus*.

## DISCUSSION

In this study, we determined the cryoEM structure of the *V. alginolyticus* lateral flagellar filament at 2.37 Å resolution (Fig. 2) and demonstrated by mass spectroscopy that the filament protein LafA is modified by pseudaminic acid at five surface-exposed serine residues in domain D1 (Figs. 3, 4). We further showed that Maf is required for lateral flagellar filament formation and motility (Figs. 5 and 6). Together, these findings provide the first structural characterization of a glycosylated *Vibrio* lateral flagellar filament and establish a direct link between flagellin glycosylation and lateral flagellar assembly.

*Salmonella* flagellins FliC and FljB consist of four domains, D0, D1, D2, and D3, arranged from the inner to the outer parts of the filament structure (26–28). The amino acid sequences of the inner D0 and D1 core domains are highly conserved among bacterial species whereas the surface-exposed D2 and D3 domains are variable (29). A notable feature of LafA is its compact architecture. Unlike *Salmonella* FliC and FljB, LafA consists only of the conserved D0 and D1 domains (Fig. 2C). Because the identified glycosylation sites are located on the surface-exposed D1 domain (Fig. 3), pseudaminic acid modification may influence interactions of the filament with surrounding environments. Recent observations that extracellular polymers such as gelatin enhance the lateral-flagellum-dependent motility of this strain raise the possibility that the filament surface properties contribute to efficient surface-associated motility (25). Future studies using site-directed mutants of the glycosylated serine residues will be required to test this hypothesis.

The conserved D0 and D1 domains are essential for filament formation and stabilization of the entire filament structure (5, 6). A major finding of this study is the identification of five pseudaminic acid modifications on LafA by the combination of cryoEM and mass spectrometry. Pseudaminic acid and related nonulosonic acids have previously been reported as flagellin modifications in diverse bacterial species, including *Campylobacter* (9, 30), *Helicobacter* (17), *Aeromonas* (31), and *Shewanella* (32). Our findings extend these observations to *Vibrio* lateral flagella and provide direct structural evidence for pseudaminic acid modification of a *Vibrio* lateral flagellin (Fig. 3). The five glycosylation sites are distributed approximately evenly over the filament surface, generating a nearly continuous glycosylated outer layer (Fig. S9). Such extensive surface modification suggests that pseudaminic acid may contribute not only to filament assembly but also to filament stability or interactions with the extracellular environment.

The close genetic linkage between *lafA* and *maf* (Fig. 1) suggested a functional relationship between the two genes, which is strongly supported by the present genetic and structural analyses. Indeed, the *maf* homolog identified downstream of *lafA* plays an essential role in lateral flagellar filament formation. Deletion of *maf* abolished motility and prevented formation of detectable lateral flagellar filaments (Fig. 5). AlphaFold3 prediction revealed that the *Vibrio* Maf homolog adopts a structure similar to previously characterized Maf proteins of *Magnetospirillum magneticum* AMB-1 (33)(Fig. S11), especially the C-terminal domain of *Vibrio* Maf and the central domain of *M. magneticum* Maf, where had exhibited similarity to sialyltransferases from *Campylobacter jejuni* (34), supporting a conserved role of Maf in flagellin modification. Nevertheless, the molecular mechanism by which Maf promotes filament assembly remains unclear. Because the flagellar type III protein export apparatus is required for efficient export and assembly of flagellar proteins (5, 6), one possibility is that glycosylation promotes efficient export and/or assembly of LafA. Alternatively, Maf-dependent modification may stabilize LafA before or after export through the flagellar protein export apparatus. Distinguishing between these possibilities will require biochemical characterization of Maf and identification of its direct substrates.

Based on the present findings and previous studies of pseudaminic acid biosynthesis, we propose a working model for lateral flagellar filament assembly in *V. alginolyticus* (Fig. 7). In this model, pseudaminic acid synthesized through the conserved biosynthetic pathway is transferred to specific serine residues of LafA by Maf prior to filament assembly. Glycosylated LafA is then exported through the flagellar protein export machinery and assembled into the lateral flagellar filament. These findings provide a structural framework for understanding how flagellin glycosylation contributes to the assembly and function of the lateral flagella in *Vibrio* species.

**Fig. 7.**
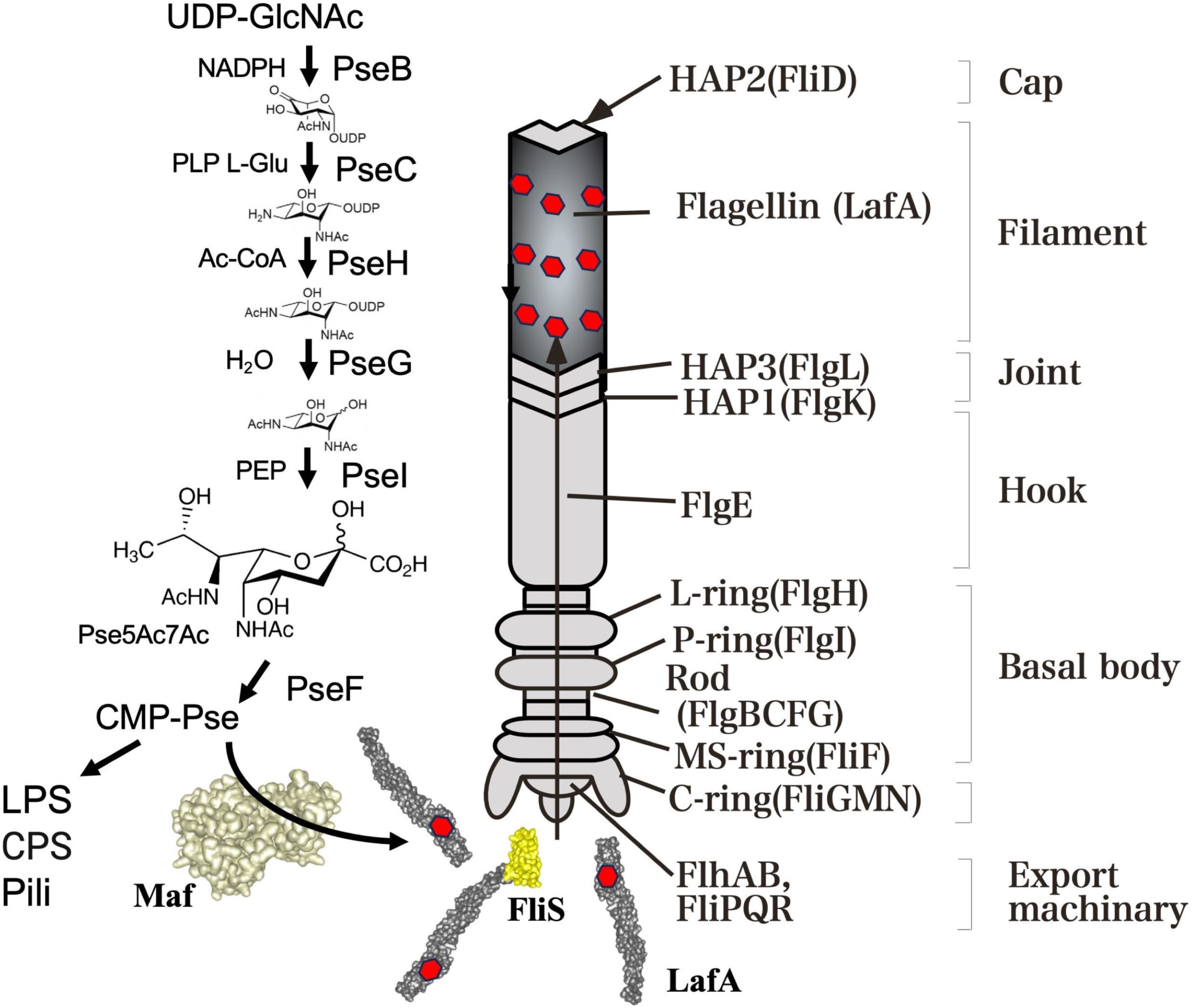
Schematic model for Maf-dependent glycosylation and assembly of *V. alginolyticus* lateral flagellar filament. Pseudaminic acid (Pse) synthesized through the conserved biosynthetic pathway is transferred to specific serine residues of LafA by Maf. Glycosylated LafA is subsequently exported by the flagellar type III protein export apparatus and assembled into the filament beneath the FliD cap. The Pse biosynthetic pathway is based on previous studies (37, 38), and the assembly model was adapted from Kint et al. (19).

## MATERIALS AND METHODS

### Strains, plasmids, and media

The bacterial strains and plasmids used here are listed in Table S1. *E. coli* cells were cultured at 37°C in LB medium [1 % (w/v) bactotryptone, 0.5 % (w/v) yeast extract, and 0.5 % (w/v) NaCl]. *V. alginolyticus* cells were cultured at 30°C in VC medium [0.5% (w/v) hipolypepton, 0.5% (w/v) yeast extract, 0.4% (w/v) K2HPO4, 3% (w/v) NaCl, and 0.2% (w/v) glucose], VPG medium [1% (w/v) hipolypepton, 0.4% (w/v) K_2_HPO_4_, 3% (w/v) NaCl, and 0.5%(w/v) glycerol], or VNG medium [1% (w/v) NZ amine, 0.4% (w/v) K_2_HPO_4_, 3% (w/v) NaCl, and 0.5% (w/v) glycerol]. When necessary, chloramphenicol was added to final concentrations of 2.5 µg/ml and of 25 µg/ml for *V. alginolyticus* and *E. coli*, respectively, after autoclaved.

### Purification of *Vibrio alginolyticus* lateral flagellar filament

*Vibrio alginolyticus* strain YM19, which produces only lateral flagella, was grown on VNG agar plates to induce swarming. Cells were harvested from the bacterial lawn and suspended in 20TN200 buffer [20 mM Tris-HCl (pH 8.0), 200 mM NaCl]. Lateral flagellar filaments were detached from the cells using a blender. Cells and large debris were removed by low-speed centrifugation, and flagellar filaments were recovered by ultracentrifugation.

The resulting filament preparation was subjected to size-exclusion chromatography using a Superose 6 10/300 column. SEC running buffer was 20TN100 buffer [20 mM Tris-HCl (pH 8.0), 100 mM NaCl]. For analysis of filament depolymerization, samples were incubated at 60°C for 10 min prior to chromatography. Eluted fractions were analyzed by SDS-PAGE followed by Coomassie Brilliant Blue staining.

Selected fractions were further characterized by LC-MS/MS, negative-stain electron microscopy, and cryo-electron microscopy. For cryoEM analysis, peak fractions containing lateral flagellar filaments were vitrified and imaged using a 300-kV transmission electron microscope.

### LC-MS/MS analysis for protein identification

For protein identification in the purified filament fraction, protein bands excised from SDS-PAGE gels were subjected to in-gel tryptic digestion. Gel pieces were washed with 50% (v/v) methanol/50 mM NH4HCO3, dehydrated with acetonitrile, dried, and incubated with trypsin (Promega, Madison, WI) at 37°C for 16 h.

Tryptic peptides were extracted with 50% (v/v) acetonitrile/0.3% (v/v) formic acid, dried under vacuum, reconstituted in 10% (v/v) acetonitrile/0.3% (v/v) formic acid, and subjected to LC-MS/MS analysis.

Tryptic peptides were analyzed using an Agilent 1290 Infinity II UPLC system coupled to a 6470 triple quadrupole mass spectrometer (Agilent Technologies). Peptide identification was performed based on MS/MS spectra against the *V. alginolyticus* protein database.

### Negative-stain electron microscopy

For observation of bacterial cells, cells were collected from agar plates and suspended in 1% (w/v) potassium phosphotungstate (pH 7.0). Samples were applied to carbon-coated copper grids and examined using a JEM-1400 transmission electron microscope (JEOL, Japan).

For observation of purified lateral flagellar filaments, amorphous carbon grids were glow-discharged using a JEC-3000FC sputter coater (JEOL, Japan). Purified filament samples (3 μL) were applied to the grids and negatively stained with 2% (w/v) uranyl acetate. Staining was repeated three times before air-drying. Images were recorded using a JEM-1400Flash transmission electron microscope (JEOL, Japan) operated at 100 kV.

### CryoEM sample preparation and data collection

Quantifoil grids (R1.2/1.3, Cu 200 mesh) were glow-discharged using a JEC-3000FC sputter coater (JEOL, Japan) at 20 mA for 20 s. A total of 2 μL of sample was applied sequentially to both sides of the grid. Grids were prepared using a Leica EM GP2 (Leica Microsystems, Germany) equilibrated at 4 °C and 100% humidity. The grids were blotted for 7 s and then immediately plunged into liquid ethane. Excess ethane was removed using filter paper, and the grids were stored in liquid nitrogen. CryoEM image datasets were acquired using SerialEM ver. 4.0.29 and yoneoLocr ver. 1.0.30 on a JEM-Z300FSC (CRYO ARM™ 300, JEOL, Japan) operated at 300 kV, equipped with a K3 direct electron detector (Gatan, Inc.) in CDS mode. The Ω-type in-column energy filter was operated with a slit width of 20 eV for zero-loss imaging. The nominal magnification was 60,000×, corresponding to a calibrated pixel size of 0.786 Å. The nominal defocus range was −0.5 to −2.0 μm. Each movie was fractionated into 60 frames (0.038 s per frame; total exposure time, 2.29 s), with a total dose of 80 e^-^/Å².

### CryoEM image processing

The gain reference was generated from 500 movies in the dataset with the “relion estimate gain” program in RELION 4.0. CryoEM data processing was performed using cryoSPARC ver. 4.6.2. A total of 6,825 movies were subjected to motion correction and CTF estimation. Micrographs with an estimated CTF resolution better than 5 Å were selected, resulting in 6,598 micrographs for further analysis.

Filament segments were initially manually picked from a subset of micrographs (44 micrographs) to generate 2D class averages for template creation. These templates were used for the first round of filament tracing, and the extracted particles were subjected to 2D classification to obtain improved class averages.

The resulting high-quality 2D classes were then used as templates for a second round of filament tracing, yielding 2,905,795 particles. These particles were subjected to two rounds of 2D classification, resulting in 1,576,907 particles.

The resulting 1,576,907 particles were subjected to ab initio reconstruction into four classes. Three classes representing similar filament structures were selected for further processing. As no significant structural differences were observed among these classes, particles from the three classes were combined for subsequent analysis.

The combined particles were subjected to helical refinement, followed by further 2D classification, ab initio reconstruction, and helical refinement. Particles were then re-extracted at a higher pixel sampling for subsequent refinement.

Duplicate particles were removed, and the remaining particles were subjected to helical refinement. Global and local CTF refinement was performed, followed by additional helical refinement. Final helical refinement using 566,405 particles yielded a reconstruction at 2.37 Å resolution, as estimated using the gold-standard FSC criterion of 0.143.

### Model building

The atomic model of the LafA filament was built using the AlphaFold-predicted structure of LafA as an initial model. The model was initially fitted into the cryo-EM map using UCSF ChimeraX (version 1.11.1) and manually adjusted in Coot (version 0.9.8.95). Real-space refinement was performed using PHENIX (version 1.20.1-4487). Model validation was carried out using MolProbity implemented in PHENIX, and this process was iterated several times. The final model contains eleven LafA subunits. CryoEM data collection, refinement, and validation statistics are summarized in Table 1.

### Glycopeptide analysis of LafA

For identification of glycosylated LafA peptides, the LafA band excised from SDS-PAGE gels was processed using an optimized in-gel digestion protocol described previously (https://doi.org/10.1002/sscp.202200121). Briefly, gel pieces were destained, reduced, alkylated, and digested overnight at 37°C with sequencing-grade modified trypsin. Peptides were recovered from the gel pieces without post-digestion acetonitrile dehydration, desalted, and subjected to LC-MS/MS analysis.

NanoLC–MS/MS analysis of digested peptides was performed using a nano-flow reverse-phase LC system (Dionex UltiMate 3000, Thermo Fisher Scientific) coupled to a Q Exactive Orbitrap mass spectrometer (Thermo Fisher Scientific, Waltham, MA, USA) via a nano-electrospray ionization source (AMR Inc., Tokyo, Japan). Desalted peptides were loaded onto a C18 reversed-phase capillary column (NTCC-360/100-3-125, 120 × 0.1 mm; Nikkyo Technos, Tokyo, Japan). Mobile phase A consisted of water containing 0.5% acetic acid, and mobile phase B consisted of 80% acetonitrile containing 0.5% acetic acid. Peptides were separated at a flow rate of 0.5 μL/min using a linear gradient of 5–40% solvent B over 100 min, followed by washing at 95% B and re-equilibration at 5% B, for a total run time of 115 min. The mass spectrometer was operated in data-dependent acquisition (DDA) mode. The ion source conditions were as follows: capillary temperature, 250 °C; spray voltage, 2.0 kV in positive ion mode; and S-lens RF level, 50. Full MS spectra were acquired over an m/z range of 350–1800 at a resolution of 70,000 with an automatic gain control (AGC) target of 3 × 10⁶. Higher-energy collisional dissociation (HCD) MS/MS spectra were acquired at a resolution of 17,500 with an AGC target of 1 × 10⁵, an isolation window of 2.0 m/z, a maximum injection time of 60 ms, and a normalized collision energy (NCE) of 27. Dynamic exclusion was set to 20 s. Raw MS/MS data were processed using Proteome Discoverer version 2.4.1.15 (Thermo Fisher Scientific). Database searches were performed using SEQUEST HT against the *Vibrio alginolyticus* protein database or the flagellin protein sequence together with the cRAP contaminant database. Search parameters included trypsin specificity with up to two missed cleavages, a precursor mass tolerance of 10 ppm, and a fragment mass tolerance of 0.02 Da. Carbamidomethylation of cysteine was set as a fixed modification, whereas oxidation of methionine, pseudaminic acid (+316.127 Da) modification of serine residues, and dHex2Pent (+410.164 Da) glycosylation were specified as variable modifications. Peptide-spectrum matches (PSMs) were filtered to a false discovery rate (FDR) of 1% using the Percolator algorithm. Modification site localization confidence was evaluated using ptmRS. Peptide identifications and glycosylation assignments were further manually inspected based on MS/MS fragmentation patterns.

### Electrospray (ESI)-TOFMS

ESI-MS was carried out using an AccuTOF JMS-T100LC mass spectrometer (JEOL, Tokyo, Japan). The peptide samples were dissolved in 50% aqueous MeOH containing 2% AcOH and loaded into a glass nanospray tip (HUMANIX Co., Hiroshima, Japan). The tip was positioned manually using an in-house built manipulator. Mass measurements were performed in positive ion mode by applying a voltage of +1200 V to the nanospray tip and 120 V between the two skimmers located behind the tip. The flow rate of the drying gas (N2) was set to 1.9 L/min, and the source temperature behind the first skimmer was maintained at 80 °C.

### Plasmid construction and mutant generation

The *lafA* and *maf* genes were amplified by PCR from genomic DNA of *V. alginolyticus* strain 138-2 and cloned into the arabinose-inducible expression vector pTSK29 for complementation analysis. DNA assembly was performed using standard cloning procedures or the NEBuilder HiFi DNA Assembly Kit (New England Biolabs).

For construction of in-frame deletion mutants, approximately 500-bp regions upstream and downstream of *lafA* or *maf* were amplified by PCR and joined to generate deletion alleles. The resulting DNA fragments were cloned into the suicide vector pSW7848, generating plasmids pTSK189 (Δ*lafA*) and pTSK190 (Δ*maf*).

Deletion mutants were constructed in *V. alginolyticus* strain YM19 by allelic exchange essentially as described previously (35, 36). Suicide plasmids were transferred from *Escherichia coli* β3914 into YM19 by conjugation. Single-crossover integrants were selected on chloramphenicol-containing plates, and double-crossover recombinants were isolated by counterselection on arabinose-containing plates. Candidate mutants were screened by colony PCR, and the deletions were confirmed by DNA sequencing.

All plasmids used in this study are listed in Table S1. DNA sequencing was performed using the BigDye Terminator v3.1 Cycle Sequencing Kit at the Nagoya University DNA Sequencing Core Facility.

### Motility assays

Motility was assessed on VNG agar plates containing either 0.3% (w/v) agar or 1.25% (w/v) agar containing 1% gelatin. Where indicated, chloramphenicol (2.5 μg mL⁻¹) and arabinose (0.1%) were added to maintain plasmids and induce gene expression. An aliquot (2 μL) of an overnight culture was spotted onto the center of each plate and incubated at 30°C in a humidified container. Colony expansion was documented by scanning the plates at the indicated time points.

### Observation of swimming behavior

*V. alginolyticus* cells were cultured overnight in VC medium at 30℃. The overnight culture was suspended in VPG medium at 50- or 100-fold dilution and incubated at 30℃ for 6h, and cells were observed under a dark-field microscope and recorded by CCD camera on computer or by a digital camera.

## Acknowledgements

We thank Kimika Maki and Yoshie Kushima for negative-stain electron microscopy. This work was partially supported by JSPS KAKENHI Grant Numbers JP24K09393 (to M.H.) and JP23K27140 (to S.K.). This work has also been supported by Research Support Project for Life Science and Drug Discovery (BINDS) from AMED under Grant Number JP23am121003, JP24am121003 and JP25am121003 (to K.N.), and by JEOL YOKOGUSHI Research Alliance Laboratories of The University of Osaka (to K.N.).

## Data Availability Statement

All data generated during this study are included in the article and supporting information. The cryoEM density map has been deposited in the Electron Microscopy Data Bank under accession code EMD-81163, and the corresponding atomic coordinates have been deposited in the Protein Data Bank under accession code 27HA.

## Author contributions

T.M., S.K., K.N., and M.H. designed the study; M.K, H.N., E.M-S., T.T., S.K., and M.H. performed the experiments; M.K., T.M., T.T., A.S., and M.H. analyzed the data; T.M., M.K., T.T., S.K., K.N., and M.H. wrote the manuscript.

