## Supplemental Figures and Table for "Structural characterization of a pseudaminic acid-modified lateral flagellar filament from *Vibrio alginolyticus*"

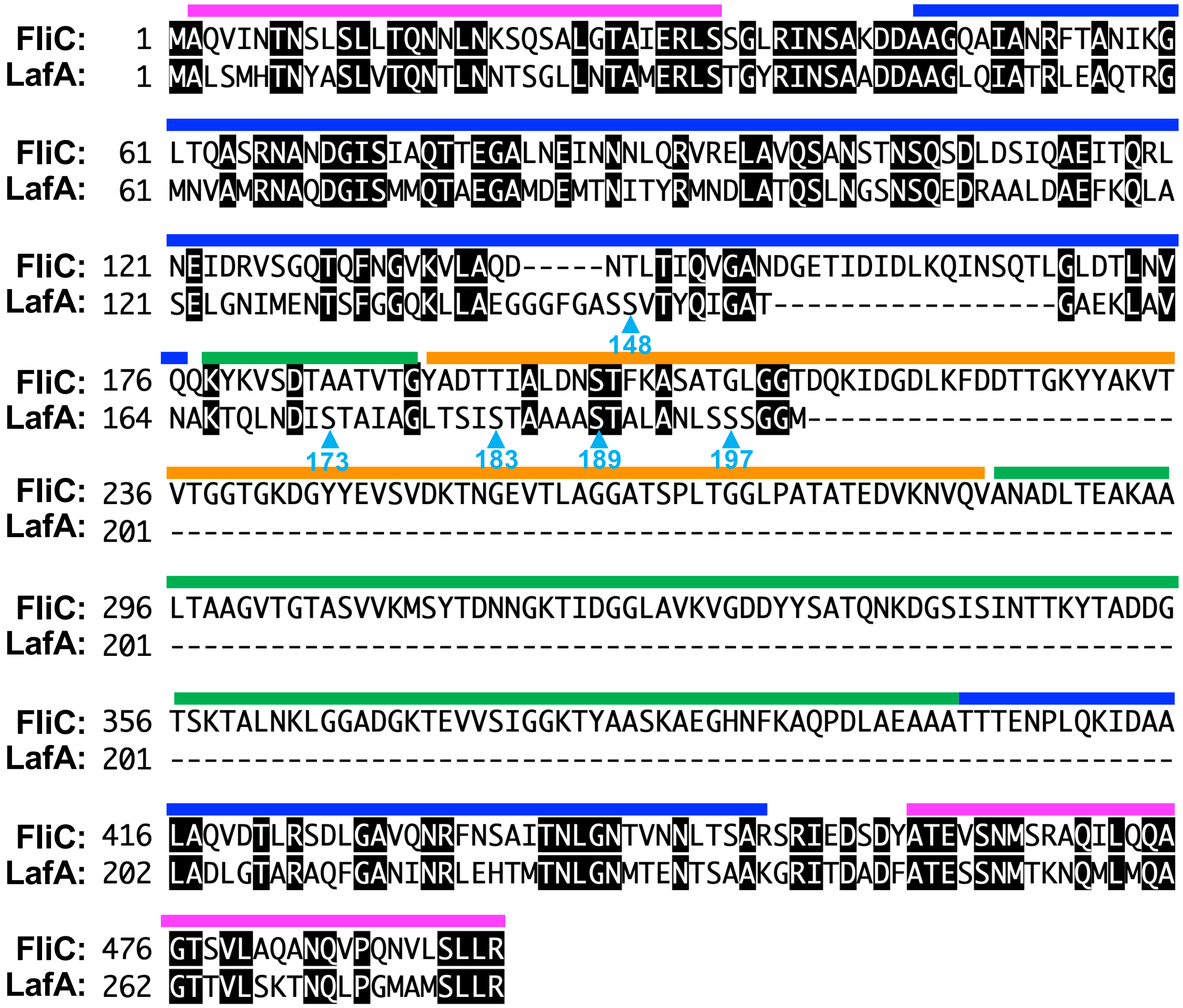

**Fig. S1 Sequence comparison of *V. alginolyticus* LafA and *Salmonella* enterica FliC.** Amino acid sequence alignment of *Salmonella* FliC (NP_460912) and *V. alginolyticus* LafA (BCB45292). Identical residues are highlighted with black boxes. Sky blue arrowheads indicate the five glycosylated serine residues identified in the LafA filament. The initiator methionine residues of both FliC and LafA are removed after protein synthesis. *Salmonella* FliC consists of four domains, D0, D1, D2, and D3, arranged from the inner to the outer parts of the filament structure. Magenta, blue, green, and orange bars indicate domains D0 (residues 2–32 and 459–494), D1 (residues 44–176 and 402–454), D2 (residues 177–189 and 284–401), and D3 (residues 190–283) of *Salmonella* FliC, respectively.

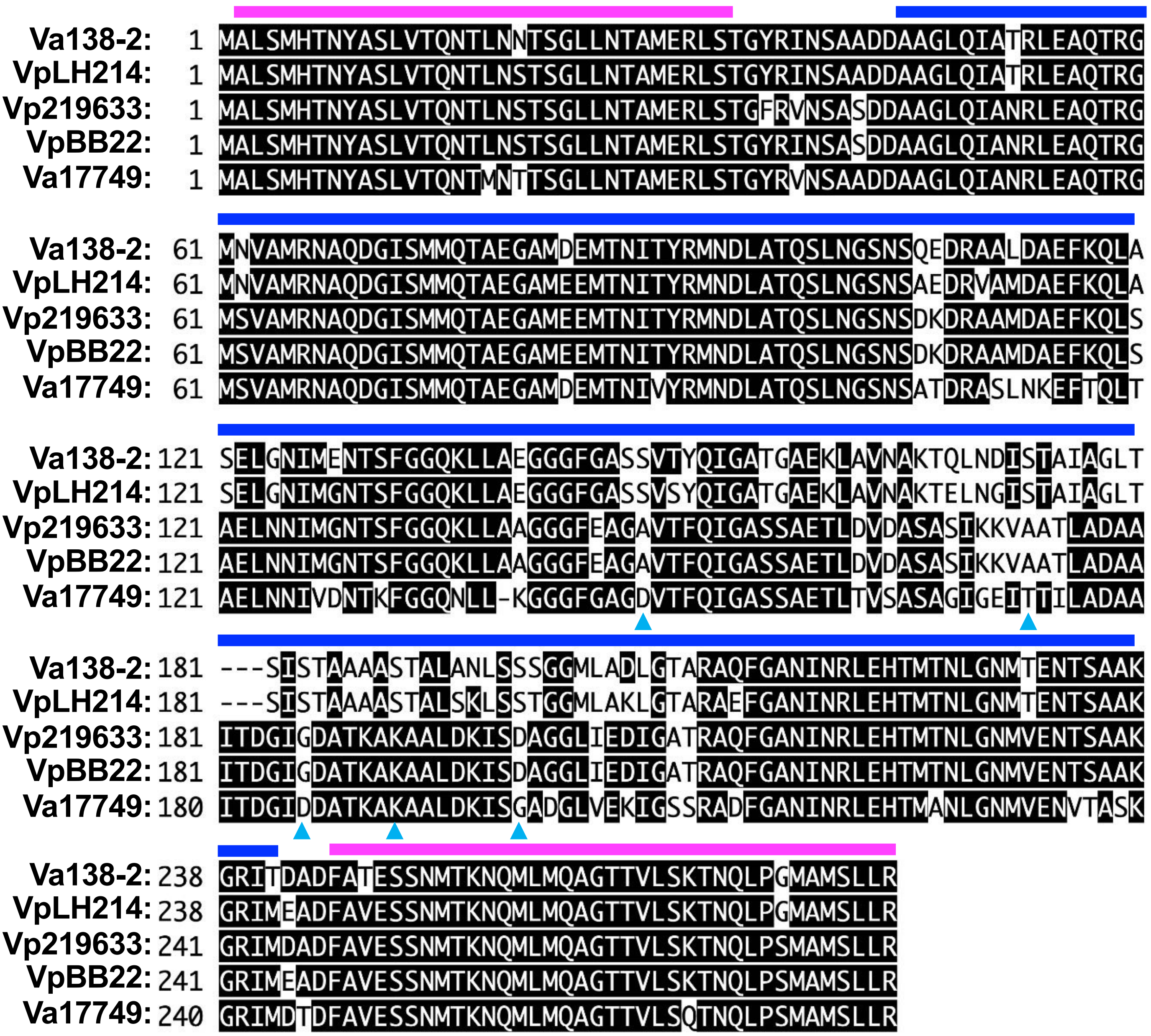

**Fig. S2 Sequence comparison of LafA homologs from *Vibrio* species.** Amino acid sequence alignment of LafA homologs from *Vibrio alginolyticus* 138-2 (Va138-2, BCB45292), *Vibrio parahaemolyticus* LH214 (VpLH214, WAG35041), *Vibrio parahaemolyticus* 2219633 (XZC27143), *Vibrio parahaemolyticus* BB22 (VpBB22, AGB12789), and *Vibrio alginolyticus* ATCC17749 (Va17749, AGV19076). Identical residues are highlighted with black boxes. Magenta and blue bars indicate domains D0 and D1 of *V. alginolyticus* LafA, respectively. LafA homologs from strains carrying a *maf* homolog exhibit sequence variations predominantly within the surface-exposed D1 domain, which contains the identified glycosylation sites as indicated by arrowheads.

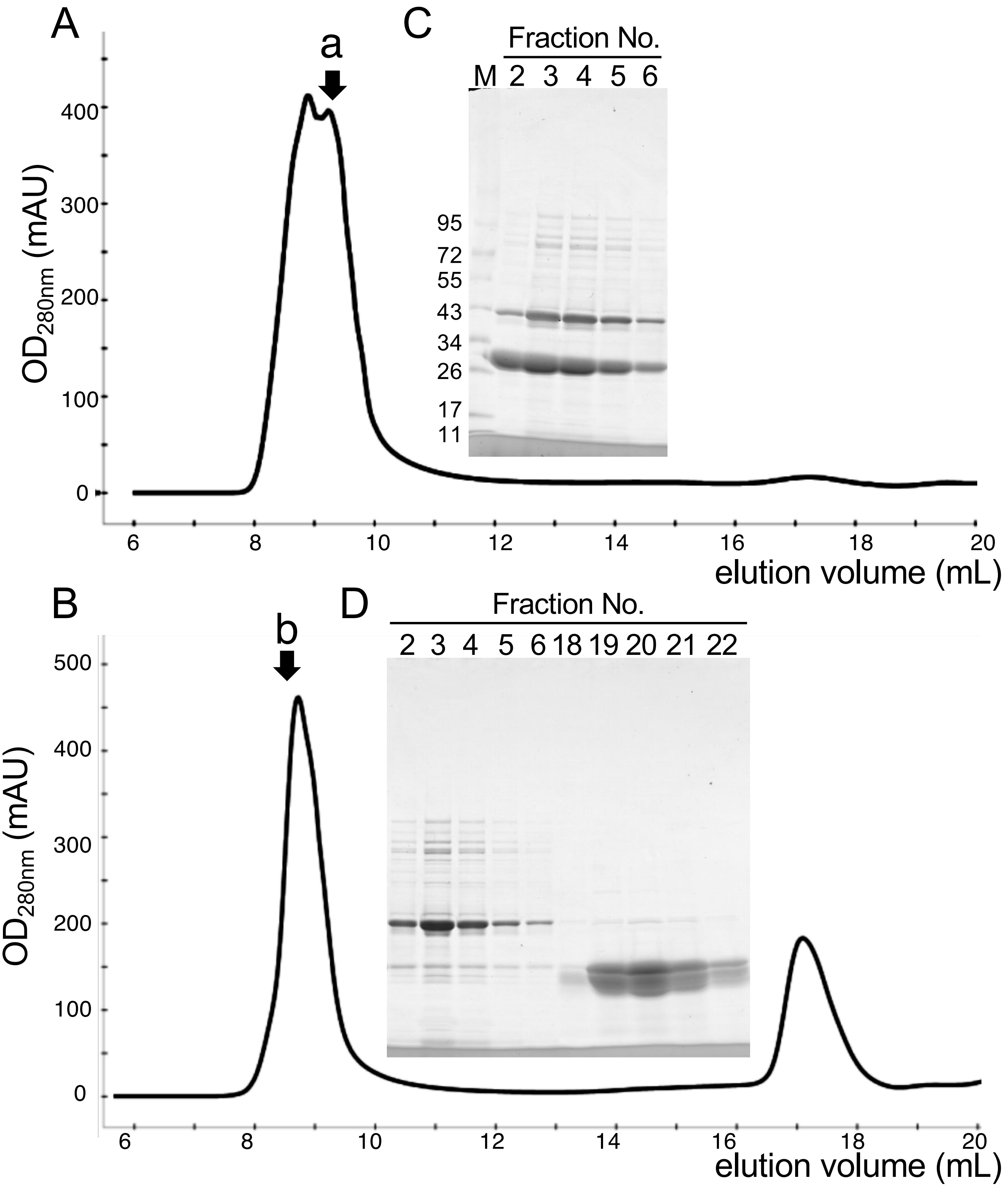

**Fig. S3 Purification of lateral flagellar filaments from *Vibrio alginolyticus* strain YM19.** The lateral flagellar filaments were isolated from YM19 cells by ultracentrifugation, suspended in 20TN200 buffer, and subjected to size-exclusion chromatography on a Superose 6 10/300 column before (A) or after heat treatment at 60° C for 10 min (B). Fractions were collected and analyzed by SDS-PAGE followed by Coomassie Brilliant Blue staining (C and D). Heat treatment induced depolymerization of the filament structure into monomeric flagellin, resulting in a shift of the elution profile toward lower molecular weight fractions. Lane M in panel C indicates molecular weight markers (kDa).

| | Gene | PEP | PSM | Coverage | MW | Sequence Coverage |
|---|---|---|---|---|---|---|
| 1. | LafA | 384 | 2665 | 67% | 29.3 | |
| 2. | OmpT | 154 | 272 | 56% | 37.5 | |
| 3. | OmpU | 74 | 87 | 45% | 35.1 | |
| 4. | FlgE | 117 | 85 | 59% | 41.1 | |
| 5. | LamB | 101 | 82 | 54% | 46.1 | |

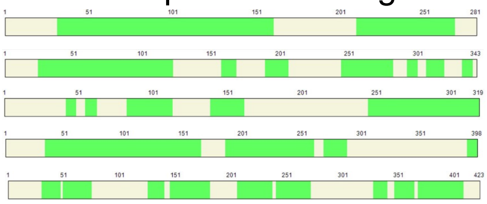

**Fig. S4 Mass spectrometric identification of proteins in the gel-filtration peak fraction.** Proteins present in the gel-filtration peak fraction (arrow a in Fig. S3A) were identified by LC-MS/MS after tryptic digestion. The table summarizes the identified proteins together with their Mascot ion scores (PEP), peptide-spectrum match counts (PSM), sequence coverage (Coverage), molecular weights (MW), and locations of the detected peptides within each protein sequence highlighted in green. The analysis confirmed the presence of LafA-derived peptides in the peak fraction.

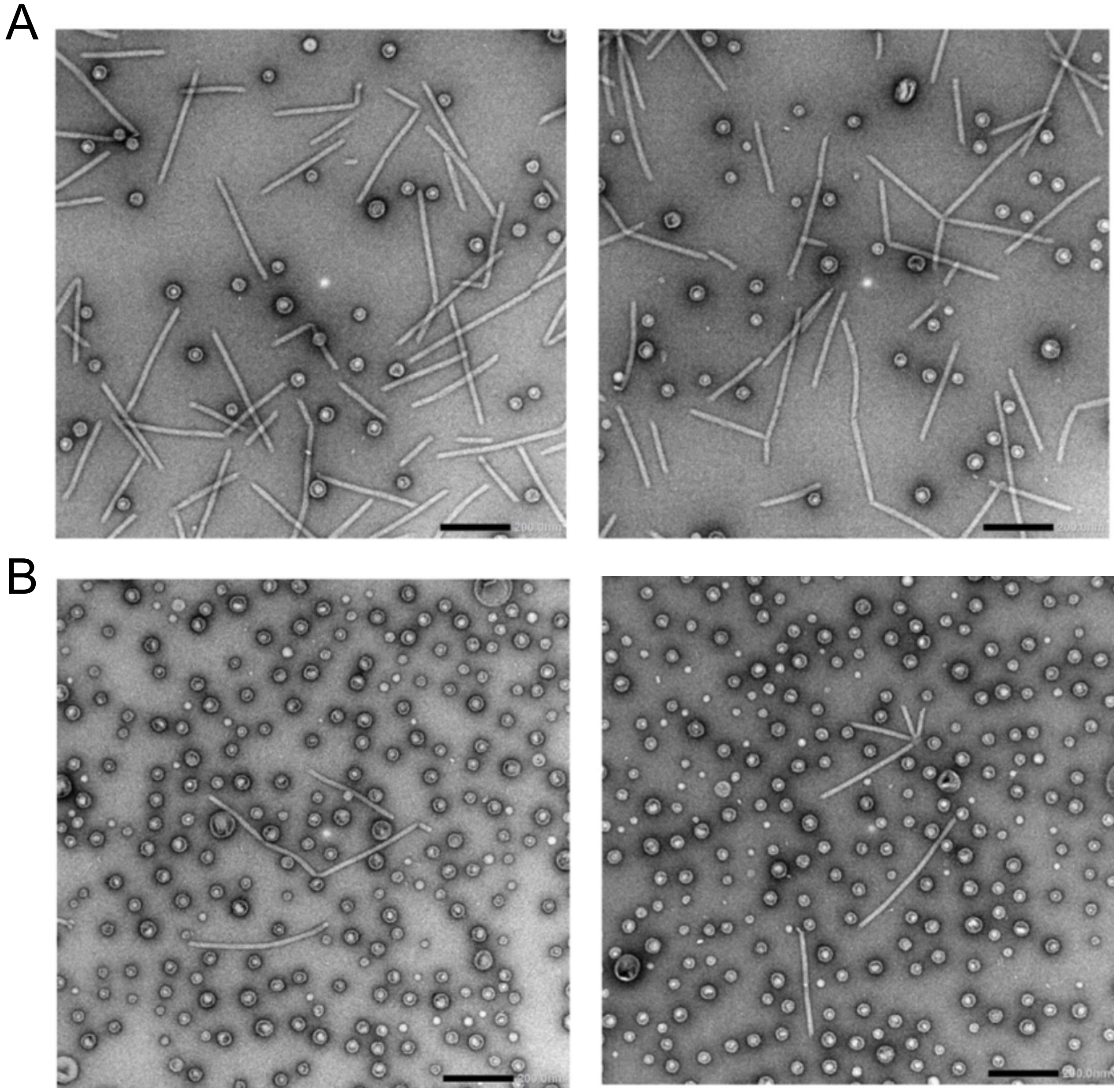

**Fig. S5 Negative-stain electron micrographs of gel-filtration fractions.** Representative negative-stain electron micrographs of the gel-filtration fractions indicated by arrows a and b in Fig. S3A and S3B, respectively. Samples were stained with uranyl acetate and observed at a nominal magnification of ×25,000. Scale bars, 200 nm. Fraction a contained numerous short filamentous structures together with vesicle-like particles, whereas fraction b obtained after heat treatment was enriched in vesicle-like particles and contained markedly fewer filamentous structures.

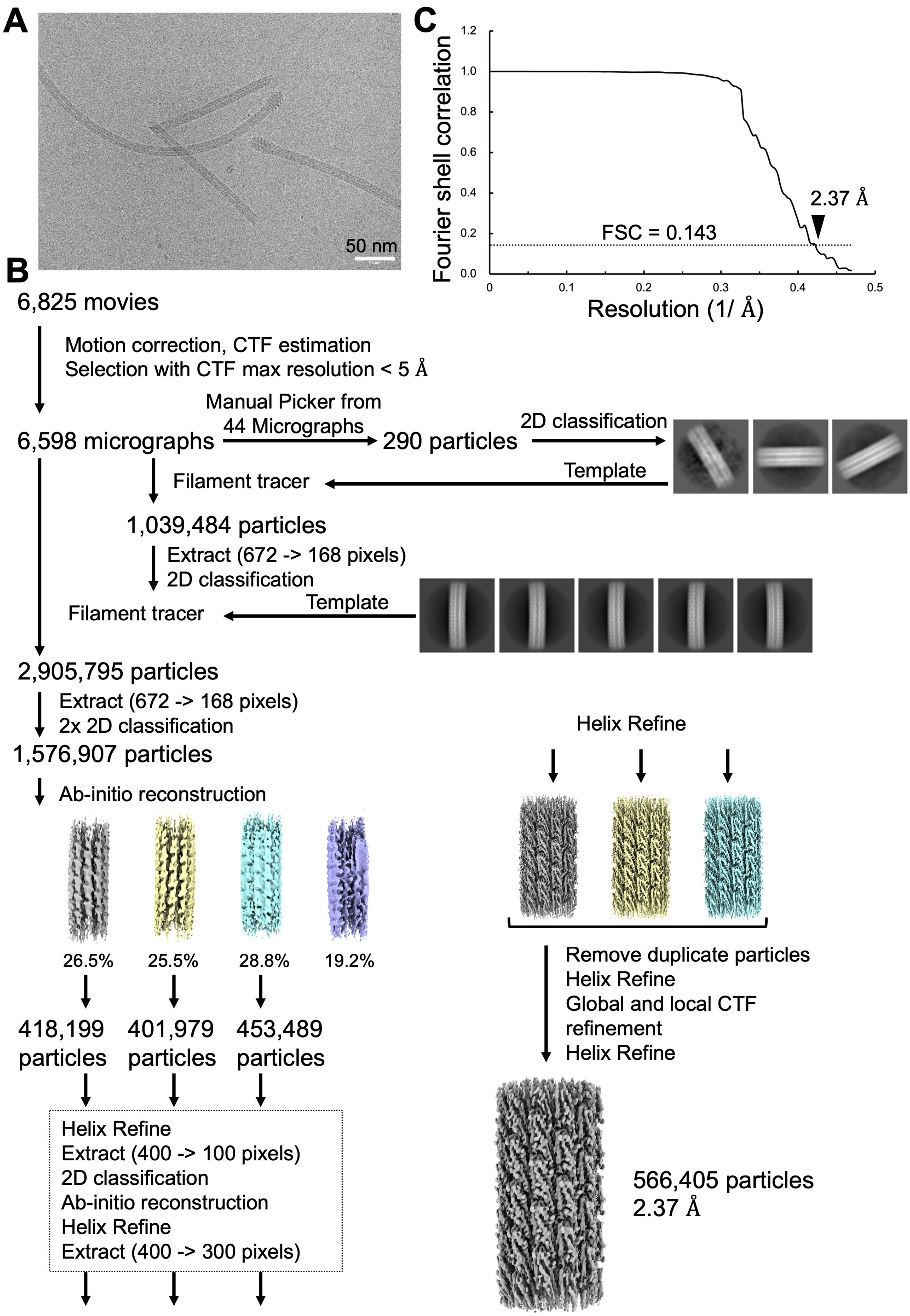

**Fig. S6 CryoEM data processing workflow of *V. alginolyticus* LafA filament.** (A) A representative cryoEM micrograph of the lateral flagellar filament. Scale bar, 50 nm. (B) CryoEM image processing workflow used for reconstruction of the LafA filament density map. (C) Fourier shell correlation (FSC) curve of the final density map of the LafA filament. The final map was reconstructed to an overall resolution of 2.37 Å (EMD-81163).

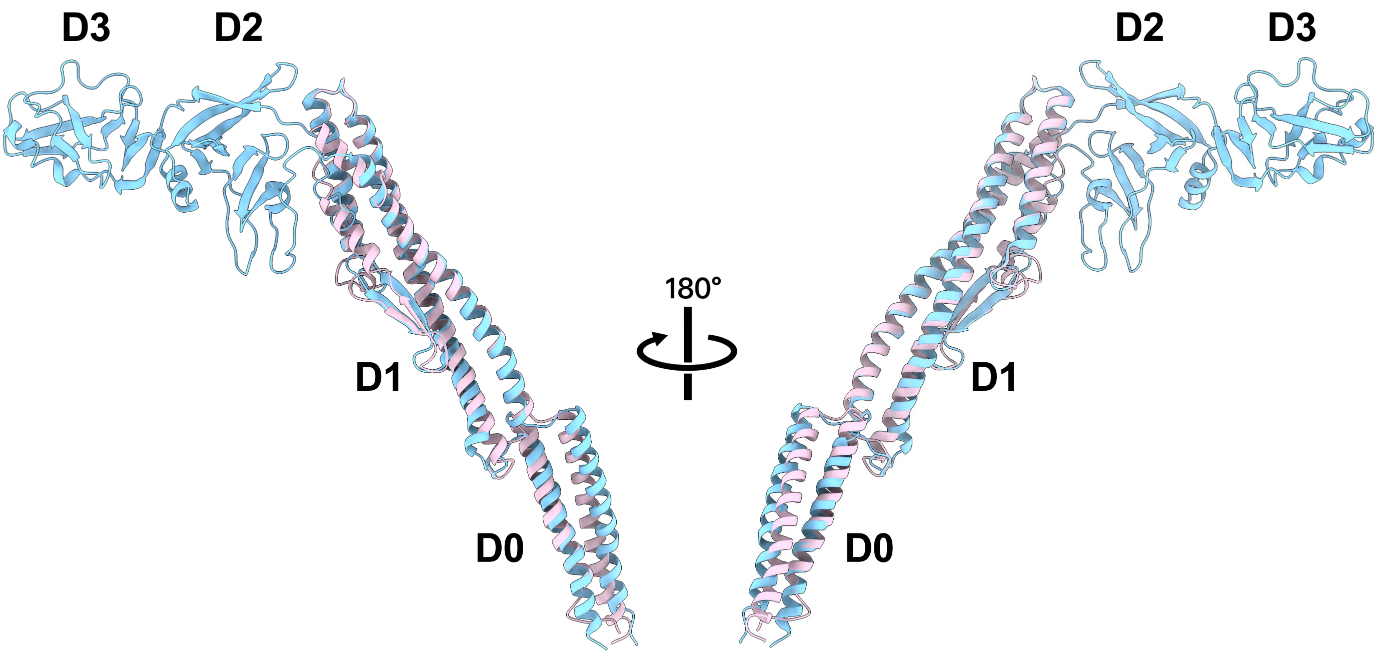

**Fig. S7 Structural comparison of *V. alginolyticus* LafA and *S. enterica* FliC.** LafA (pale mauve, PDB ID: 27HA) is superimposed onto FliC (sky blue, PDB ID: 1UCU) with an RMSD of 0.936 Å. While the structures of conserved D0 and D1 domains are highly similar between the two proteins, LafA lacks the large surface-exposed D2 and D3 domains characteristic of FliC, resulting in a more compact and thinner filament architecture.

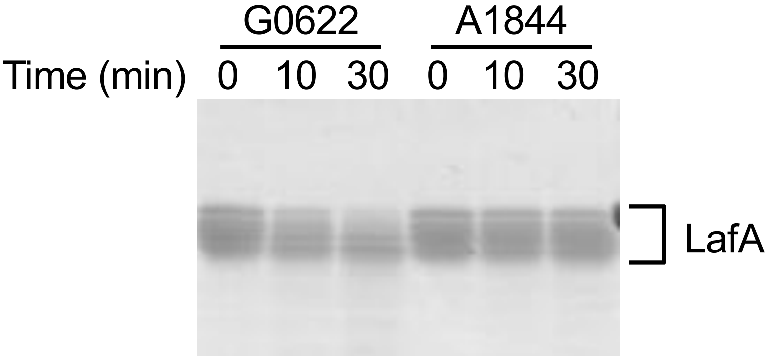

**Fig. S8 Glycosidase treatment of lateral flagellin.** Monomeric LafA prepared from heat-depolymerized lateral flagellar filaments was treated with the glycosidases indicated above the lanes at 37ºC. Samples collected at the indicated time points were analyzed by SDS-PAGE and visualized by CBB staining. G0622, Endo-α-N-acetylgalactosaminidase plus sialidase; A1844, Endo-α-N-acetylgalactosaminidase alone.

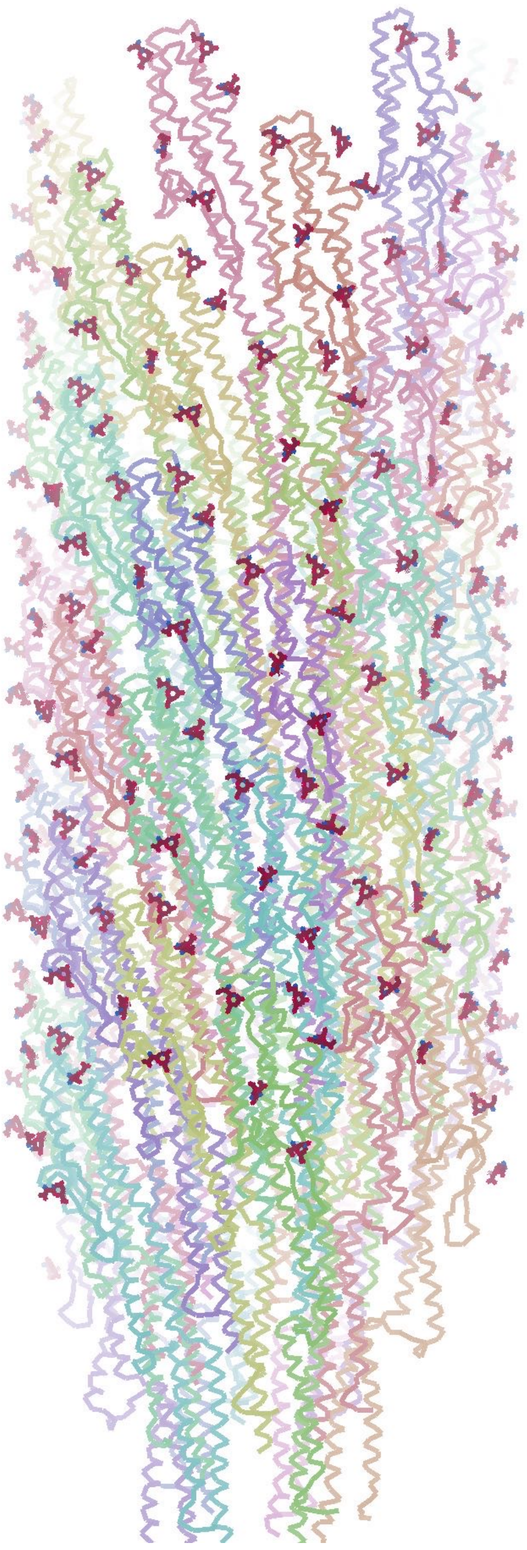

**Fig. S9 Distribution of pseudaminic acid modifications on the lateral flagellar filament.** Side view of the LafA filament model showing the distribution of pseudaminic acid (Pse) modifications on the filament surface. Pse molecules are colored red. The five glycosylation sites of each subunit are distributed approximately evenly over the filament surface, generating a nearly continuous glycosylated outer surface.

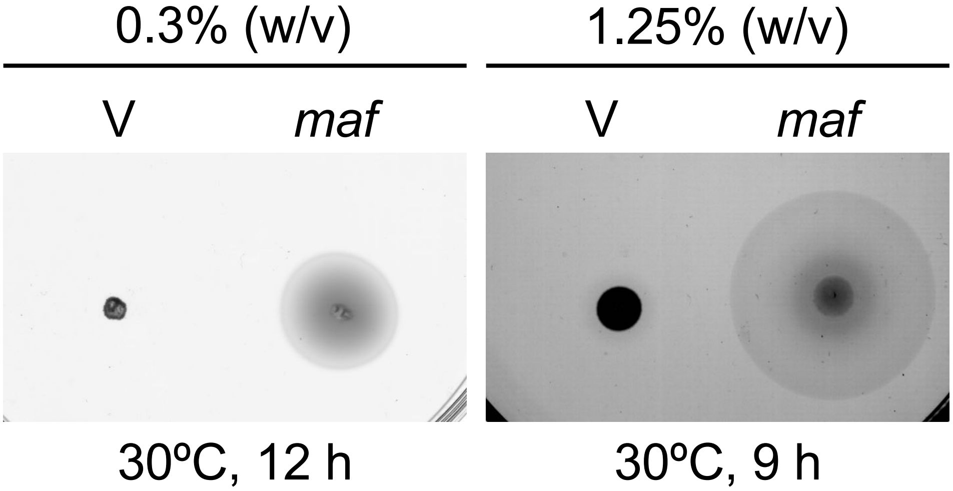

**Fig. S10 Complementation of the Δ*maf* mutant.** Representative swimming and swarming motility assays of the Δ*maf* strain carrying either the empty vector (pBAD33, V) or plasmids expressing Maf (pTSK193). Cells were inoculated onto VNG plates containing 0.3% agar (left) or 1.25% agar (right) supplemented with 1% gelatin, chloramphenicol, and arabinose. The plates were incubated at 30ºC and photographed after 12 h (left) and 9 h (right), respectively. Expression of *maf* restored lateral flagella-dependent motility of the Δ*maf* mutant.

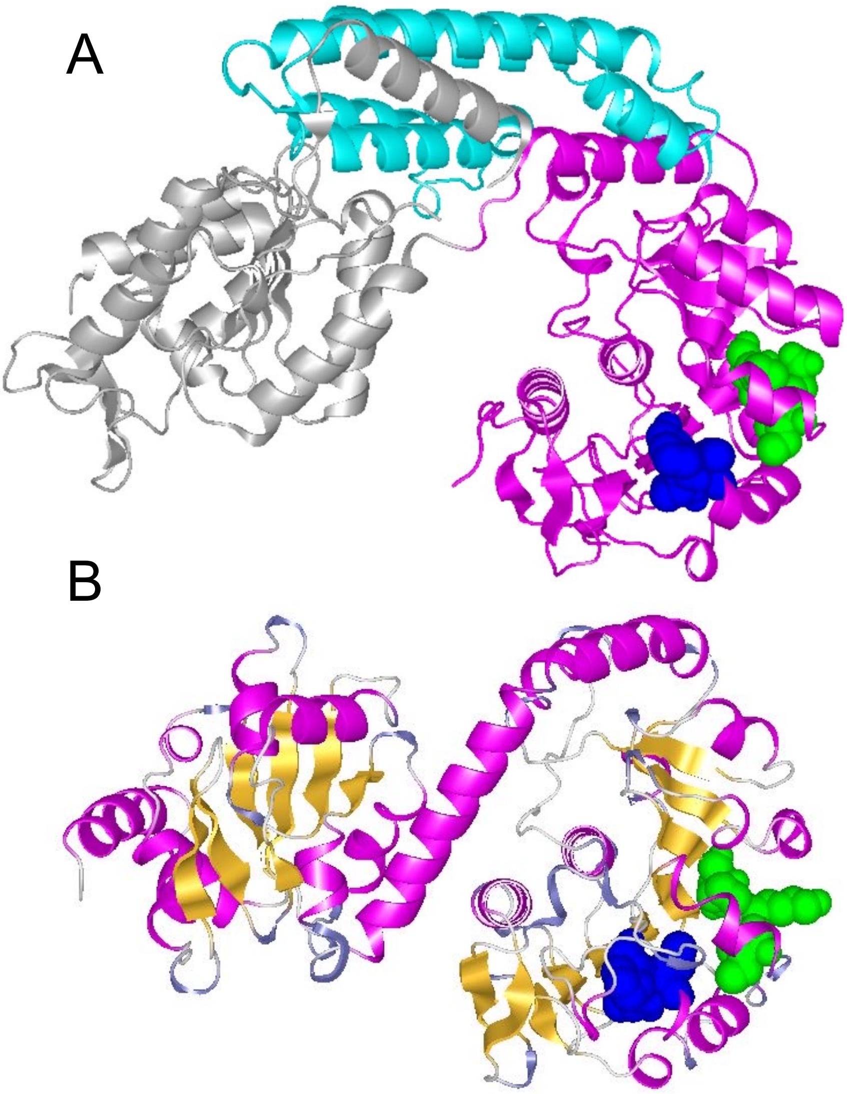

**Fig. S11 Structural comparison of Maf proteins.** (A) Crystal structure of the Maf glycosyltransferase from *Magnetospirillum magneticum* AMB-1 (PDB ID: 5MU5). The N-terminal, central, and C-terminal domains are colored silver, purple, and light blue, respectively. (B) AlphaFold3-predicted structure of the *V. alginolyticus* Maf homolog, colored as secondary structure. The predicted structure of Maf exhibits similarity to the C-tarminal domain of Maf and the middle domain of *M. magneticum*. Conserved GPSL and IKPD motifs for glycosyltransferase are shown as space-filling models in blue and yellow-green, respectively.

**Table S1. Strains and plasmids used in this study.**

| Strain or plasmid | Genotype or description | Reference or source |
|---|---|---|
| *V. alginolyticus* | | |
| 138-2 | Wild-type strain ($Pof^+$ $Laf^+$) | (1) |
| YM19 | 138-2 $Pof^-$ ($Pof^-$ $Laf^+$) | (1) |
| NMB384 | YM19 Δ*lafA* ($Pof^-$ $Laf^-$) | this study |
| NMB385 | YM19 Δ*maf* ($Pof^-$ $Laf^-$) | this study |
| *E. coli* | | |
| DH5α | Recipient for DNA manipulation | (2) |
| β3914 | Recipient for conjugational transfer of pSW7848 | (3) |
| Plasmids | | |
| pGEM-T easy | TA cloning vector, $Amp^r$ | Promega |
| pBAD33 | $Cm^r$, $P_{BAD}$ | (4) |
| pSW7848 | Suicide plasmid for allele exchange $oriV_{R6K}$ $oriT_{RP4}$ | (5) |
| pTSK29 | $Cm^r$, $P_{BAD}$, MCS with NdeI site | (6) |
| pTSK188 | The 3.2 kb PCR product (*lafA-maf* locus) in pGEM-T easy | this study |
| pTSK189_1 | The 1.0 kb PCR product flanking 500 bp up- or down-stream of *lafA* (Δ*lafA* fragment) in pGEM-T easy | this study |
| pTSK189 | Δ*lafA* fragment in pSW7848 | this study |
| pTSK190 | The 1.0 kb PCR product flanking 500 bp up- or down-stream of *maf* (Δ*maf* fragment) in pSW7848 | this study |
| pTSK191 | *lafA* in pTSK29 | this study |
| pTSK192 | *maf* in pTSK29 | this study |

$Amp^r$, ampicillin-resistant; $Cm^r$, chloramphenicol-resistant; $Pof^+$, possessing a polar flagellum; $Laf^-$, lack of lateral flagella

1. I. Kawagishi, Y. Maekawa, T. Atsumi, M. Homma, Y. Imae, Isolation of the polar and lateral flagellum-defective mutants in *Vibrio alginolyticus* and identification of their flagellar driving energy sources. *J Bacteriol* **177**, 5158-5160 (1995).
2. S. G. Grant, J. Jessee, F. R. Bloom, D. Hanahan, Differential plasmid rescue from transgenic mouse DNAs into *Escherichia coli* methylation-restriction mutants. *Proc Natl Acad Sci USA* **87**, 4645-4649 (1990).
3. F. Le Roux, J. Binesse, D. Saulnier, D. Mazel, Construction of a Vibrio splendidus mutant lacking the metalloprotease gene vsm by use of a novel counterselectable suicide vector. *Appl Environ Microbiol* 73, 777-784 (2007).
4. L. M. Guzman, D. Belin, M. J. Carson, J. Beckwith, Tight regulation, modulation, and high-level expression by vectors containing the arabinose pBAD promoter. *J Bacteriol* **177**, 4121-4130 (1995).
5. M. E. Val, O. Skovgaard, M. Ducos-Galand, M. J. Bland, D. Mazel, Genome engineering in Vibrio cholerae: a feasible approach to address biological issues. *PLoS genetics* 8, e1002472 (2012).
6. S. Inaba, T. Nishigaki, N. Takekawa, S. Kojima, M. Homma, Localization and domain characterization of the SflA regulator of flagellar formation in *Vibrio alginolyticus. Genes Cells* 22, 619-627 (2017).
